# Arachidonic acid metabolism controls macrophage alternative activation through regulating oxidative phosphorylation in PPARG dependent manner

**DOI:** 10.1101/2020.10.15.340265

**Authors:** Miao Xu, Xiaohong Wang, Xudong Jia, Yongning Li, Xue Geng, Lishi Zhang, Hui Yang

**Author notes:** Corresponding Author: Lishi Zhang., Address: No.16, Section 3, Renmin South Road, Chengdu 610041, China, Hui Yang, Address: No.7 Panjiayuan Nanli, Beijing 100021, China.

## Abstract

Macrophages polarization is mainly controlled by metabolic reprogramming in microenvironment, thus leading to distinct outcomes of various diseases. However, the role of lipid metabolism in the regulation of macrophage alternative activation is incompletely understood. Using human THP-1 and mouse bone marrow derived macrophages polarization models, we revealed a pivotal role for arachidonic acid metabolism in controlling the polarization of M1 and M2 macrophages. We demonstrated that M2 macrophage polarization was inhibited by arachidonic acid, but inversely facilitated by its derived metabolite prostaglandin E2 (PGE2). Furthermore, PPARG bridges these two unconnected processes via modulating oxidative phosphorylation. These results highlight the critical role of arachidonic acid metabolism as an immune regulator in modulating metabolic homeostasis and pathological process.

## Introduction

Many diseases including obesity, cardiovascular diseases and tumor are tightly associated with energy metabolism, which provides fuel for these physiological or pathological processes. Energy metabolic homeostasis profoundly impact immune responses in tissue microenvironment (Patel et al., 2019). When energy was surplus, immune cells reprogrammed their metabolic pathway to trigger metaflammation (Brestoff and Artis, 2015). Obesity is a prototypical example of how energy metabolic homeostasis affect immunological function. Lipids depositing in various tissues leads to hypoxia and adipocyte stress thus recruits innate immune cells and promotes chronic activation of survival pathway (Koelwyn et al., 2020). In return, phenotype change of immune cells can function to regulate system or local metabolic state (Odegaard and Chawla, 2013).

Macrophage as one of the prominent components of immune system is versatile. They adopt different polarization state depending on the context. Macrophage sensors, integrates and responses to stimulus from its local microenvironment to regulate metabolic tissues homeostasis through inflammation or insulin action (Odegaard and Chawla, 2011). Metabolic reprogramming of tumor microenvironment (TME) by cancer cells provide a perfect example of how metabolism interplay with immune function (Pavlova and Thompson, 2016). Cancer cells released lactate, glutamine, succinate and α-ketoglutarate (α-KG) help T cell and macrophage polarize towards immunosuppressive phenotype (Angelin et al., 2017, Chen et al., 2017, Mehla and Singh, 2019). In contrast, metabolic reprogramming of the TAM inhibited tumor progression by allowing the accumulation of T cell receptor engineered T cells (Stromnes et al., 2019). Dysfunction of macrophages often contribute to systemic inflammation, thus maintaining of the normal state of macrophage is critical for health state (Hotamisligil, 2017). Based on functional diversity, macrophages are mainly divided into two phenotypes, classically activated macrophages (M1) and alternatively activated macrophages (M2). The process from resting macrophages to M1 or M2 macrophages is called macrophage polarization. The differentiation and polarization of macrophage is intimately related to energy metabolism of the microenvironment. Metabolic homeostasis especially within adipose and liver tissues has been found closely related to M2 macrophages, which can promote insulin sensitivity (Odegaard and Chawla, 2011, Biswas and Mantovani, 2012). However, whether the metabolic alternation is instructive or responsive during function changes of macrophages is still unknown. Emerging evidences reveal that macrophage use glucose or fatty acid as fuel source to impact its differential activation (Saha et al., 2017). How these energy metabolism especially lipid metabolism contribute to macrophage polarization remains unclear.

In this study, we aim to explore the mechanism of macrophage polarization associated with lipid metabolism. Combining analysis of transcriptome profiles and lipid metabolomics, we report that arachidonic acid metabolism determines polarization of M1 and M2 macrophages. Arachidonic acid (AA) and its key metabolite prostaglandin E2 (PGE2) directly regulate macrophages polarization in IL-4/IL-13 induced M2 macrophages. However, PPARG activation by specific agonist rosiglitazone strikingly reverses PGE2 induced M2 polarization. PGE2 enhances oxidative phosphorylation (OXPHOS) through modulating PPARG during M2 polarization. Our data implicate arachidonic acid metabolism as an intrinsic regulator of M2 macrophage polarization----by affecting OXPHOS---that is essential for optimal polarization.

## Results

### Lipid metabolism closely correlates to macrophage polarization

To explain how lipid metabolism function in the regulation of macrophage polarization, we constructed THP-1 derived macrophage polarization model and validated basic features including morphology, surface markers, mRNA and cytokines(Fig S1). To profile metabolic signature of induced macrophages, we analyzed transcriptome profiles of M (LPS+IFNγ) and M (IL4+IL13), which in this manuscript we simply wrote as M1, M2, respectively. As Fig 1 and Fig S2 suggested, two phenotype of macrophages had divergent energy metabolism features compared with static M0 macrophages. Activation of genes controlling fatty acid biosynthesis (FAS) (Fig 1A) and oxidative phosphorylation (OXPHOS) (Fig 1D) were enhanced in M2 macrophages. Thus, to investigate whether these pathways contribute to M2 activation, series of targeted inhibitors were applied to this polarization model. As Fig 1B, 1C and 1E suggested, FAS inhibition by FASN-IN-4 tosylate (FAI), Fatostatin (FATO), FT113 (FT) dose-dependently reduced M2 polarization. Similarly, OXPHOS inhibition by 3-Nitropropanoic acid (NP), VLX600 (VLX) and IACS-10759 (IA) (Fig 1F-H) could dose-dependently attenuate M2 macrophage polarization. Other lipid utilization associated processes including lipolysis, fatty acid transport (FAT) and fatty acid oxidation (FAO) showed more complex alternation with both up and down regulated genes (Fig S2A-C). However, inhibition of these processes by particular inhibitors revealed polarization impacts as well (Fig S2D-F). As Fig S2D showed, FAT inhibition as well as FAO inhibition significantly inhibited M2 polarization, which in line with FAS and OXPHOS inhibition. The exception to this trend occurred with lipolysis inhibition, which dramatically promote M2 macrophage activation. These data suggest that up-regulation of fatty acid biosynthesis and utilization were crucial for M2 macrophage polarization.

**Figure 1.**
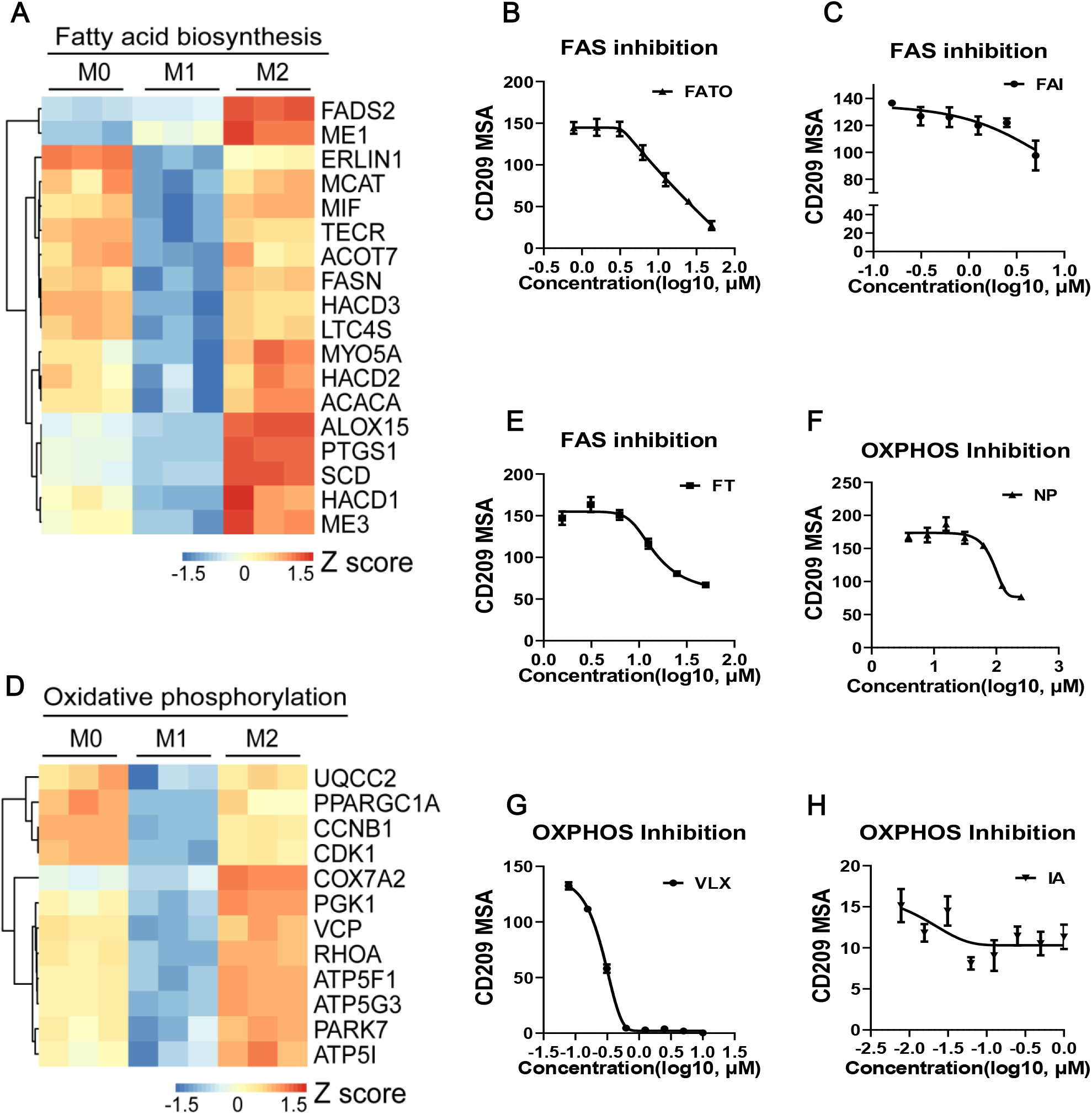
Lipid metabolism closely correlates to macrophage polarization. (A and D) Gene expression in THP-1 derived differentially activated macrophage related to fatty acid biosynthesis or oxidative phosphorylation, respectively. (B, C and E) CD209 expression curves of THP-1 derived M2 macrophages via high content screening system (HCSS) with specific fatty acid biosynthesis (FAS) inhibitors treated as indicated for 48 hours. (F-H) CD209 expression curves of THP-1 derived M2 macrophages via HCSS with specific oxidative phosphorylation (OXPHOS) inhibitors treated as indicated for 48 hours. Error bars represent the mean ± SEM from 3 biological replicates. MSA, mean stain area.

### Arachidonic acid metabolism is a hallmark of M2 macrophage

Given that lipid metabolic characteristics closely related to macrophage polarization, we questioned which one contribute most to M2 macrophage polarization. To test this, we integrated profiles of transcriptomics and metabolomics to analyze determinants. All differentially expressed genes (DEGs) between M1 and M2 were displayed in Fig 2A, in which 1645 genes were up-regulated in M2. Computational overlapping of all identified genes with the Molecular Signatures Database (MSigDB; Broad Institute) hallmark gene sets suggested an enrichment of arachidonic acid metabolism in M2 macrophages (Fig 2B). Relative abundance of genes matching arachidonic acid metabolism were significantly higher in M2 macrophages (Fig 2C). Heightened expression of these genes was largely related to prostaglandins and leukotrienes production. In accordance, the key metabolic enzymes for AA, 15-lipoxygenase (15-LO, encoded by ALOX15) and cyclooxygenases (COX-1/COX-2, encoded by PTGS1/PTGS2) were significantly induced in M2 macrophages (Fig 2D).

**Figure 2.**
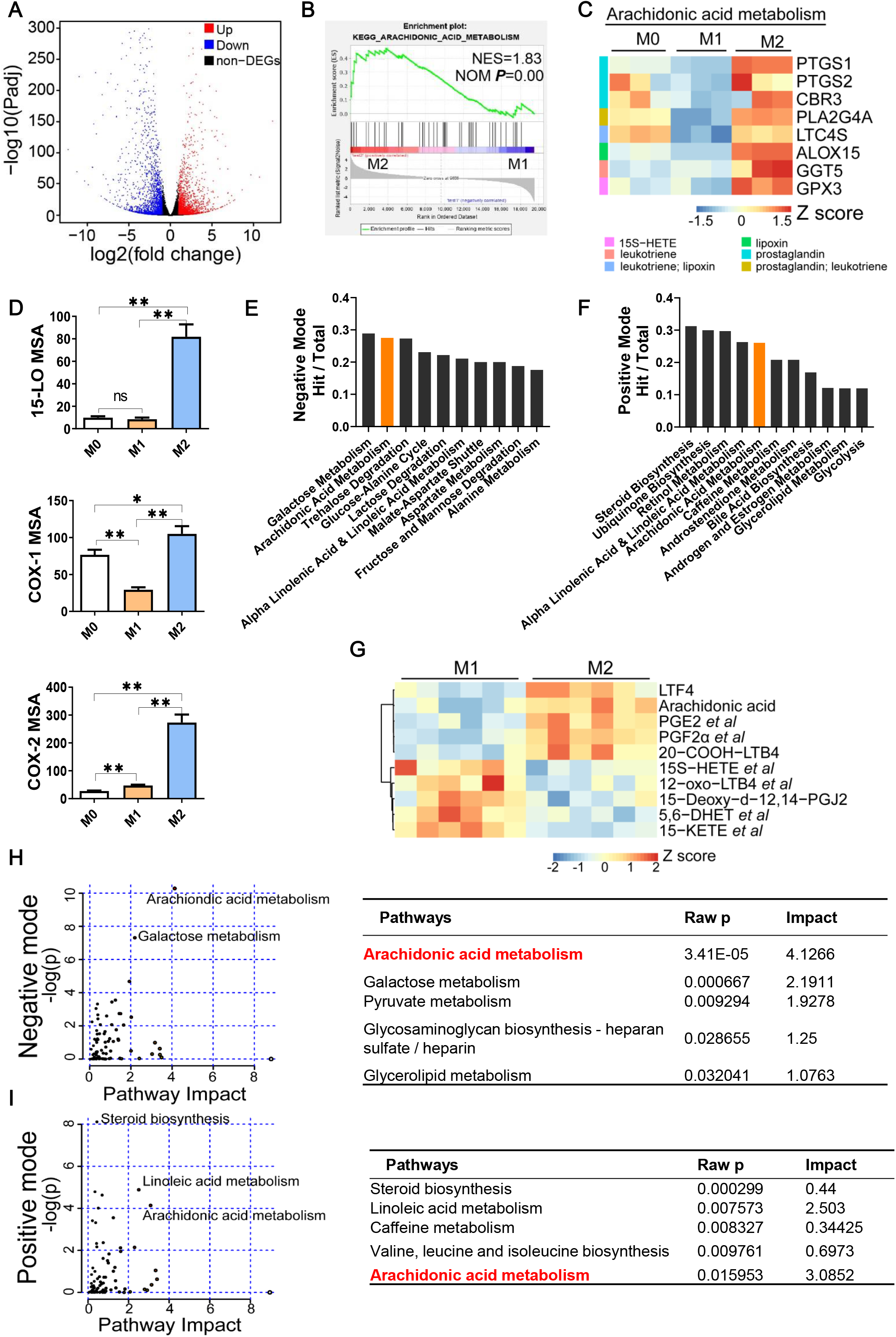
Arachidonic acid metabolism is a hallmark of M2 macrophages. (A) Differentially expressed genes (DEGs) in M2 macrophages; fold changes are in comparison with M1 macrophages. Genes with |log2(fold change)| ≥ 1 and Padj ≤ 0.05 were seen as DEGs. (B) Enrichment plot for arachidonic acid metabolism in THP-1 derived M2 macrophages from Gene Set Enrichment Analysis (GSEA). (C) Heatmap of DEGs matching “arachidonic acid metabolism” expression signature according to KEGG Pathway Analysis of RNA-seq data from M0,M1,M2 macrophages with three biological replicates. (D) Protein expression of key metabolic enzymes for arachidonic acid via HCSS in M0,M1,M2 macrophages. ✱P< 0.05; ✱✱ P < 0.01. Error bars represent the mean ± SEM from three biological replicates. (E and F) Top 10 pathways of negative or positive ion mode from Metabolites Set Enrichment Analysis (MSEA). (G) Heatmap of differentially expressed metabolites matching “arachidonic acid metabolism” expression signature according to KEGG Pathway Analysis of lipidomics data from M1,M2 macrophages with six biological replicates. (H and I) Enrichment pathways from integrated transcriptomics and lipidomics data by Joint Pathway Analysis. Left panel: Enrichment plots. Right panel: corresponding information for left plots.

To better understand the lipid metabolites signature during macrophage polarization, we analyzed metabolomics of M1 and M2 macrophages. PCA and PLS-DA analysis revealed distinguished clusters of M1 and M2 macrophages (Fig S3A, S3B). A total of 3652 and 2328 differential ions (identified as 808 and 510 differential metabolites) were obtained in positive and negative mode, respectively (Fig S3C). We next performed Metabolites Set Enrichment Analysis (MSEA) to these differential metabolites. Top 10 enriched pathways were displayed in Fig 2E, 2F. Arachidonic acid metabolism was the only two pathways that included in both modes, ranking 2^nd^ and 5^th^ respectively. Another included pathway, alpha linoleic acid and linoleic acid metabolism, shares some key enzymes with arachidonic acid metabolism such as 15-LO which could explain its enrichment. Next, we compared relative abundance of differential metabolites in arachidonic acid metabolism pathway. As shown in Fig 2G (details in Supplementary Table S2), metabolites profile of M1 and M2 were deviating and AA, some prostaglandins (PGE2, PGF2α *et al*) and leukotrienes (LTF4, 20-COOH-LTB4) were higher in M2 macrophages. We then integrated transcriptomics with lipid metabolomics by joint pathway analysis on MetaboAnalyst website. As expected, arachidonic acid metabolism was the most remarkable pathway with highest pathway impact in both modes (Fig 2H, 2I). Other metabolic pathways such as linoleic acid pathway and galactose metabolism were also significantly changed but with lower pathway impact (Fig 2H, 2I). Together, these data revealed that arachidonic acid metabolism was the most remarkable lipid metabolism characteristic between M1 and M2 macrophages. Enhanced arachidonic acid metabolism could be a hallmark of M2 macrophages..

### Arachidonic acid and PGE2 inversely regulate M2 polarization

Given that M2 macrophages own enhanced arachidonic acid metabolism, we questioned whether and how it affect M2 polarization. First we directly measured polarization effect of AA by combining treatment with IL-4 and IL-13. Both surface marker (CD209 for THP-1 model, CD206 for BMDM model) and functional cytokines (IL-4, TARC) had been decreased by AA, indicating suppressed M2 macrophage polarization (Fig 3A–3D). The key enzymes for arachidonic acid metabolism are generally constitutively expressed and determines what eicosanoids a cell can synthesize. Our data (Fig 2) revealed relative higher abundance of lipoxygenases and cyclooxygenases in M2 macrophage, thus we verified how these enzymes link to polarization. As Fig S4A, S4B displayed, inhibition of lipoxygenases (by PD146176 and MK886) decreased M2 polarization in a dose-dependent manner. Consistently, inhibition of cyclooxygenases by indomethacin (INDO) decreased M2 polarization indicated by lower expression of IL-4, TARC and CD209, CD206 (Fig 3E–3H). This suggested that metabolites of AA may favor M2 polarization. Thus we assessed polarization in the presence or absence of corresponding metabolites from AA. As expected, presence of PGE2 significantly promoted M2 polarization as suggested by increasingly expressed M2 markers (IL-1RA, CD209, CD206) (Fig 3I–3L). Collectively, these data indicated a critical role for AA and its metabolite PGE2 in optimal M2 activation of macrophages in response to IL-4/IL-13.

**Figure 3.**
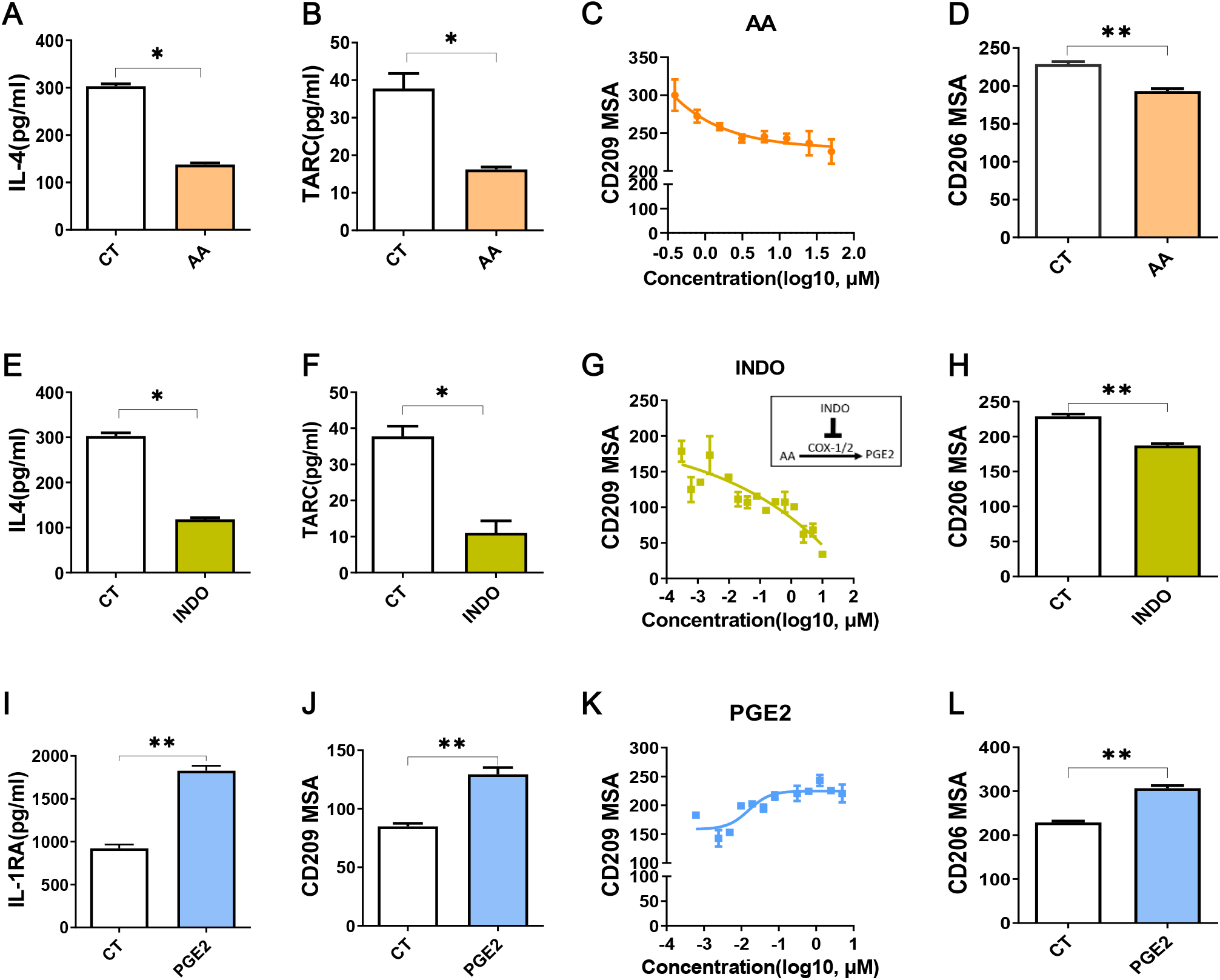
Arachidonic acid and PGE2 inversely regulate M2 polarization. (A, B, E, F and I) Cytokines of THP-1 derived M2 macrophages treated with compounds for 48h during polarization as indicated (C, G and K) CD209 expression curves of THP-1 derived M2 macrophages with treatment as indicated via HCSS. (D, H and L) Analysis of CD206 expression of BMDM derived M2 macrophages via HCSS with treatments as indicated. (J) Analysis of CD209 for THP-1 derived M2 macrophages treated as indicated during polarization via HCSS. CT represents corresponding solvent control. Arachidonic acid (AA, 50μM), indomethacin (INDO, 10μM), prostaglandin E2 (PGE2, 10μM). Error bars represent the mean ± SEM. Data presented are from three biological replicates. ✱P< 0.05; ✱✱ P < 0.01.

### PGE2 inhibits PPARG resulting in M2 macrophage polarization

Previous studies had demonstrated that PPARG were essential for M2 polarization (Odegaard and Chawla, 2011), therefore we speculated that PPARG may involve in the molecular mechanism of macrophage polarization induced by AA and PGE2. When treated with specific agonist for PPARG, rosiglitazone (R), M2 marker (CD209) was dramatically decreased dose-dependently (Fig 4A). In contrast, CD209 was significantly enhanced by the inverse agonist T0070907 (T) dose-dependently (Fig 4A), suggesting that human macrophage M2 polarization might be closely associated with PPARG de-activation. Functional cytokines secreted by M2 macrophages (IL-1RA and TARC) were correspondingly reduced by PPARG activation and TARC was increased by PPARG de-activation (Fig 4B). In consistent with THP-1 derived macrophages, PPARG activation by rosiglitazone inhibited BMDM derived macrophage M2 polarization as well (Fig 4C). These data suggested that PPARG de-activation was critical for M2 polarization.

**Figure 4.**
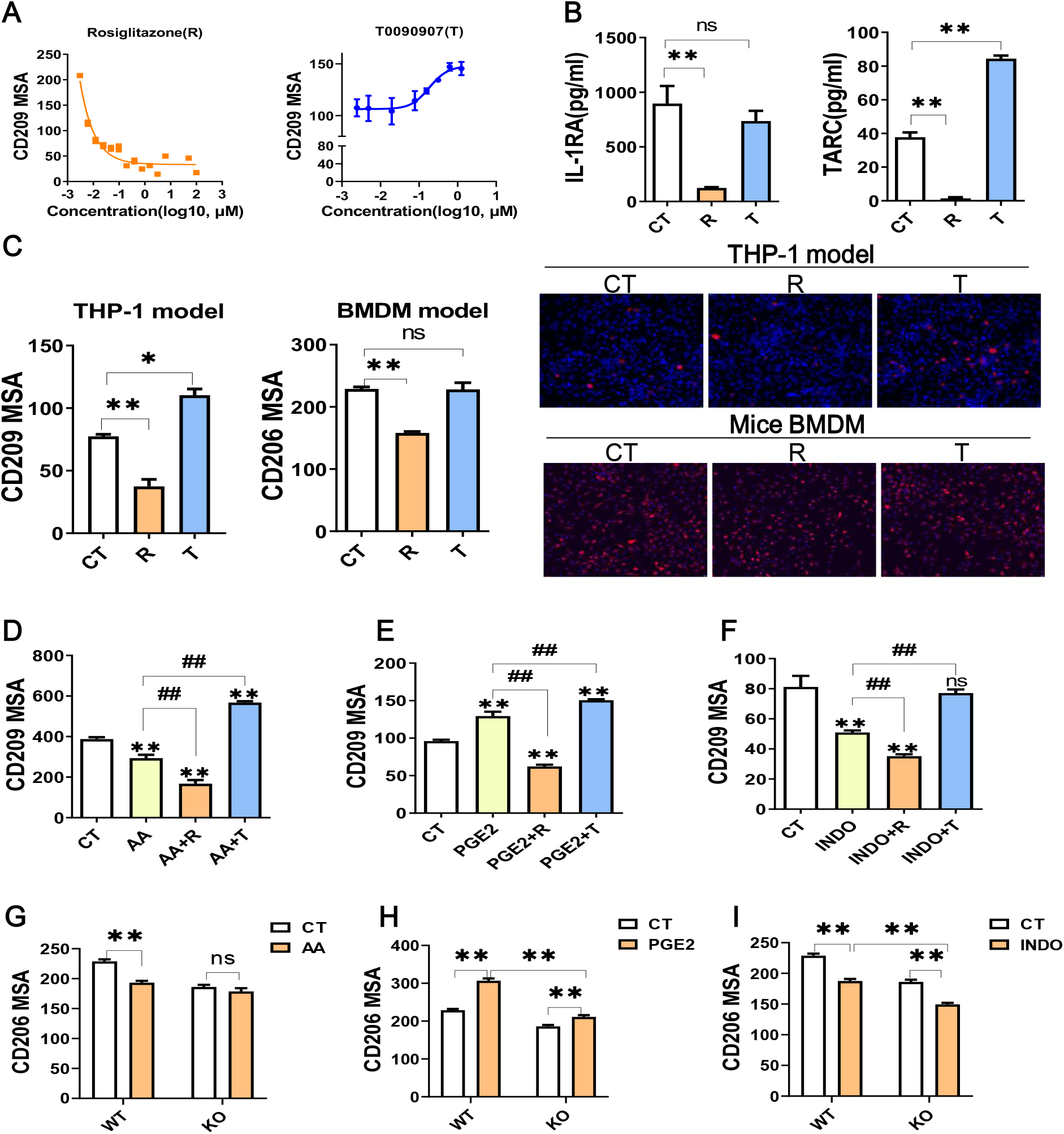
PGE2 inhibits PPARG resulting in M2 macrophage polarization. (A) CD209 expression curves of THP-1 derived M2 macrophages with treatments as indicated. (B) Cytokines of THP-1 derived M2 macrophages with treatment as indicated. (C) Protein expression of M2 markers (CD209 or CD206) from THP-1 or BMDM derived M2 macrophages with treatment as indicated. Right panel: representative images. Blue staining: nuclei; Red staining: CD209 or CD206. (D-F) Protein expression of CD209 in THP-1 derived M2 macrophages with treatment as indicated. (G-I) Protein expression of CD206 for BMDM derived M2 macrophages from wide type (WT) or monocyte specific PPARG knockout mice (KO) with treatment as indicated. CT represents corresponding solvent control. R, rosiglitazone; T, T0070907. R(10μM), T(1μM),AA(50μM), INDO(10μM), PGE2(2μM). Error bars represent the mean ± SEM from three biological replicates. Compared with CT, ✱P< 0.05; *✱ P < 0.01. ns, no significance. Compared with first treatment as indicated, ## P < 0.01.

Since inhibition of M2 polarization by AA (Fig 3A–3D) was similar to R, we presumed that AA inhibited M2 macrophage polarization through activating PPARG. To test this, we examined polarization effect of AA in the presence of T0070907. When PPARG de-activated by T0070907, AA could not inhibit M2 polarization while PPARG activated by R could enhance the inhibition of AA on CD209 expression, suggesting that the effect of AA could be reversed by PPARG de-activation while enhanced by PPARG activation (Fig 4D). At the same time, we questioned whether PGE2 promoted M2 polarization by suppressing PPARG activation. We found that PPARG activation totally reversed M2 polarization mediated by PGE2 while PPARG de-activation further enhanced M2 polarization mediated by PGE2 (Fig 4E). Reducing PGE2 production by INDO inhibited M2 polarization thus INDO shared a similar PPARG activation response with AA (Fig 4F). To formally address the possibility of PPARG bridging AA/PGE2 mediated M2 macrophage polarization, we isolated bone marrow derived monocytes (BMDM) from wild type (WT, Pparg^+/+^) or monocyte specific PPARG knockout (KO, Pparg^-/-ΔMono^) mice to construct polarization model. In WT model, M2 marker CD206 was significantly suppressed by AA while increased by PGE2 as expected. However, compared with WT, monocyte specific PPARG knockout significantly abolished or dampened these effects on M2 polarization (Fig4G, 4H), indicating a PPARG-dependent mechanism of AA/PGE2 mediated M2 polarization. Intriguingly, suppressed CD206 expression by INDO was not abolished but was enhanced by monocyte specific PPARG knockout, suggesting that INDO might have additional mechanisms besides PPARG activation to regulate M2 polarization of macrophages (Fig 4I). Together with our findings from human THP-1 derived macrophage polarization and mice BMDM polarization, these data supported the proposal of a role for PPARG bridging AA or PGE2 mediated M2 macrophage polarization.

### PGE2 enhances OXPHOS through modulating PPARG during M2 polarization

On the basis of OXPHOS was enhanced in M2 macrophages (Fig 1C) and FAO fuel OXPHOS with acetyl-CoA (Pearce and Pearce, 2013), we proposed that the observed promoting or inhibiting polarization effects of PGE2 or AA might attribute to these two processes. To test this speculation, we conducted transcriptomics analysis on M2 macrophages with PGE2 treated during polarization. Firstly, we explored the top 20 enriched pathways (ranked by normalized enrichment score, NES) in PGE2 treated cells by GSEA (Fig 5A). When considering false discovery rate q-value (FDR q value, usually no more than 0.25 was acceptable), the 11^th^ pathway, oxidative phosphorylation, was the most significantly enriched pathway with highest NES as shown in Fig 5A and supplementary table S3. The enrichment plot of OXPHOS was displayed in Fig 5B, suggesting that PGE2 treated during M2 polarization up-regulated OXPHOS remarkably. Further comparing the DEGs within this pathway demonstrated that genes related to mitochondria respiratory complex I (NDUFA8 *et al*), II (SDHA), III (UQCRH *et al*), IV(COX7A2 *et al*), V (ATP5MG *et al*) were increased (Fig 5C). To explore whether PPARG involves in the enhancement of OXPHOS by PGE2, we tried to block this effect by co treating with R. As expected, PGE2 dramatically increased the expression of ATP5A, one subunit of ATP synthase (complex V), while co treating with R weaken this effect significantly, suggesting that PPARG activation might dampen the enhancement effect of PGE2 on OXPHOS (Fig 5D). In addition, given that FAO can serve as a replenishment pathway for OXPHOS and PPARs are key regulators for FAO, we next determine the involvement of PPARG on FAO. As shown in Fig 5E, CPT1A, a key enzyme for FAO, was significantly inhibited by PPARG activation (resembling specific inhibitor ETO), while was increased by PPARG de-activation, indicating that PPARG activation inhibited FAO, thus might weaken OXPHOS. This was consist with a previous study that demonstrated inhibition of OXPHOS by PPARG activation (Lee et al., 2017). Collectively, these data revealed that PGE2 enhanced OXPHOS during M2 polarization and PPARG de-activation participated in this process.

**Figure 5.**
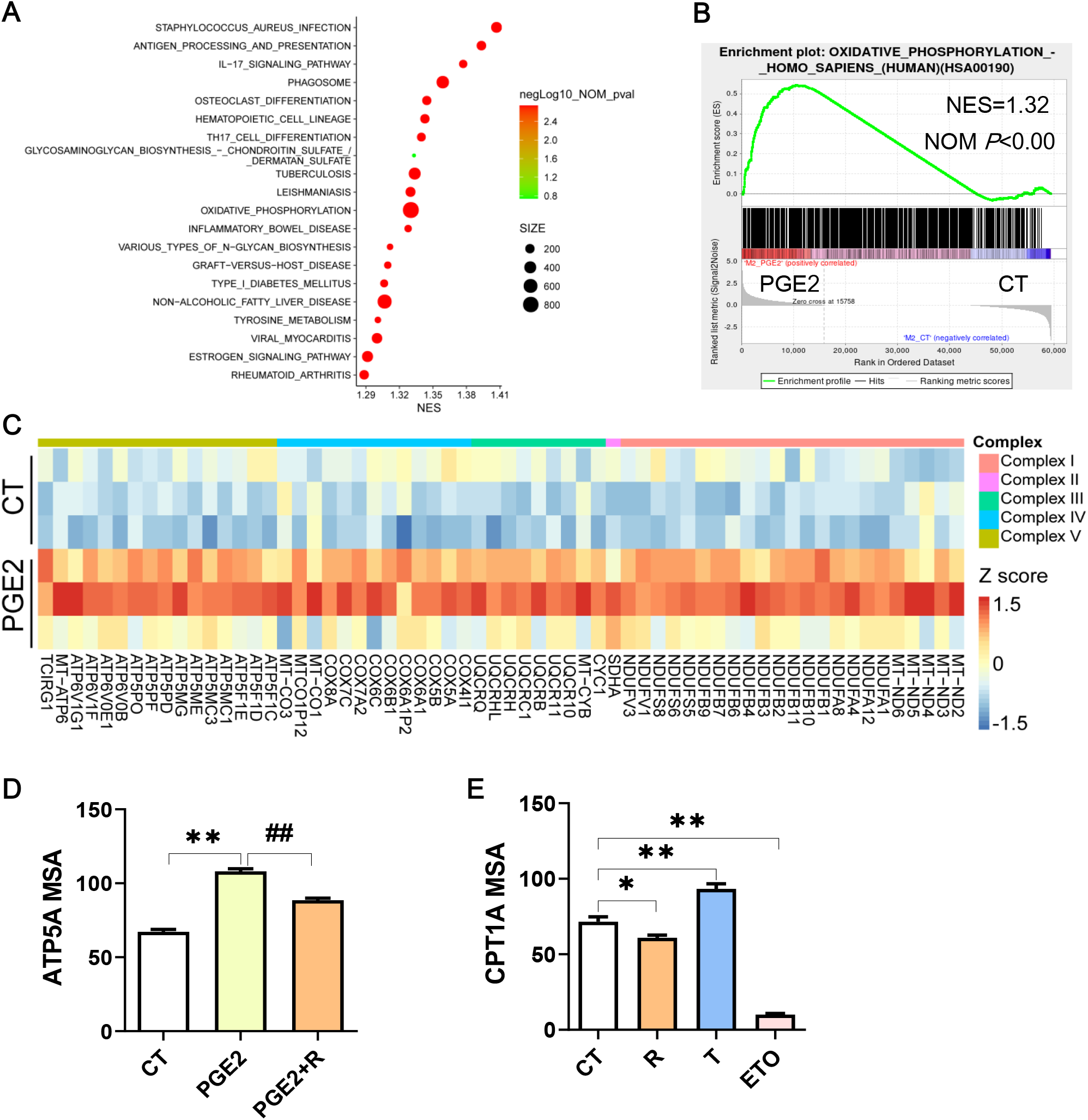
PGE2 enhances OXPHOS through modulating PPARG during M2 polarization. (A) Enrichment pathways (TOP 20, ranked with normalized enrichment score (NES)) in THP-1 derived M2 macrophages with PGE2 treated during polarization. (B) Enrichment plot for oxidative phosphorylation in THP-1 derived M2 macrophages with PGE2 treated during polarization from GSEA analysis. (C) Heatmap for representative DEGs matching “oxidative phosphorylation” of (B). (D) Protein expression of ATP5A with treatment during polarization as indicated via HCSS. (E) Protein expression of CPT1A with treatment during polarization as indicated. PGE2(2μM), R(10μM), T(1 μM), ETO (Etomoxir, 100μM, serve as positive control for CPT1A inhibition).Compared with control, ** P < 0.01. Compared with first treatment as indicated, ## P < 0.01.

### Arachidonic acid metabolism is correlated with M2 polarization in promoting tumor progression

Previous studies examining the crosstalk between cancer cells and macrophages suggested that nutrients availability may play a role in immunosuppressive tumor microenvironment (Mora et al., 2019, Gupta et al., 2017). M2 type tumor associated macrophages (M2 TAMs) formation have been seen as results of tumor cell “re-education” (Vitale et al., 2019), which means that tumor cells initiative macrophage to polarize toward pro-tumor phenotype through metabolic reprogramming, suggesting that tumor cells derived metabolites may have a role for M2 TAM formation. In our previous study, we have validated M2 TAM infiltration in esophageal carcinogenesis (Yang et al., 2020), thus we questioned whether arachidonic acid metabolism facilitate M2 TAM polarization in esophageal cancer. Unsurprisingly, in mice esophageal squamous cell carcinoma (ESCC), comparing with non-tumor tissue, several key enzymes for AA metabolism were up-regulated in tumor and M2 macrophages marker Arg1 was also increased in tumor tissues (Fig 6A), indicating a correlation between AA metabolism and M2 TAM formation. Simultaneously, using transcriptomics profiles from human ESCC, we check whether OXPHOS correlated with M2 TAM. Several FAO or OXPHOS associated genes (PPARGC1A, COX7A1, SDHA) positively correlated to M2 TAM markers (CD200R1, MRC1, CD209, CD163) as Fig 6B and Fig S5A-S5B showed, indicating the existence of energy utilization correlating to M2 TAM formation. As many markers for M2 macrophage such as MRC1, CD209, CD163 and TREM2 were positively correlated to PGE2 biosynthesis (suggested by PTGES and PTGS1) (Fig 6C), we next verified the correlation between PGE2 biosynthesis and OXPHOS. As expected, PGE2 biosynthesis and OXPHOS also displayed positive correlation (Fig 6D). This was consistent with our in vitro observation and support our hypothesis (arachidonic acid metabolism facilitate M2 TAM polarization in esophageal cancer) well. In addition, by calculating the correlation between PTGS1/PTGS2 and PPARG, we found that PGE2 biosynthesis was negatively correlated to PPARG (Fig 6E), which indirectly verified the hypothesis that PGE2 suppressed PPARG. Together, these findings remind us that AA metabolism might make contribution to M2 TAM formation via PPARG-OXPHOS modulation, thus promote tumor progression.

**Figure 6.**
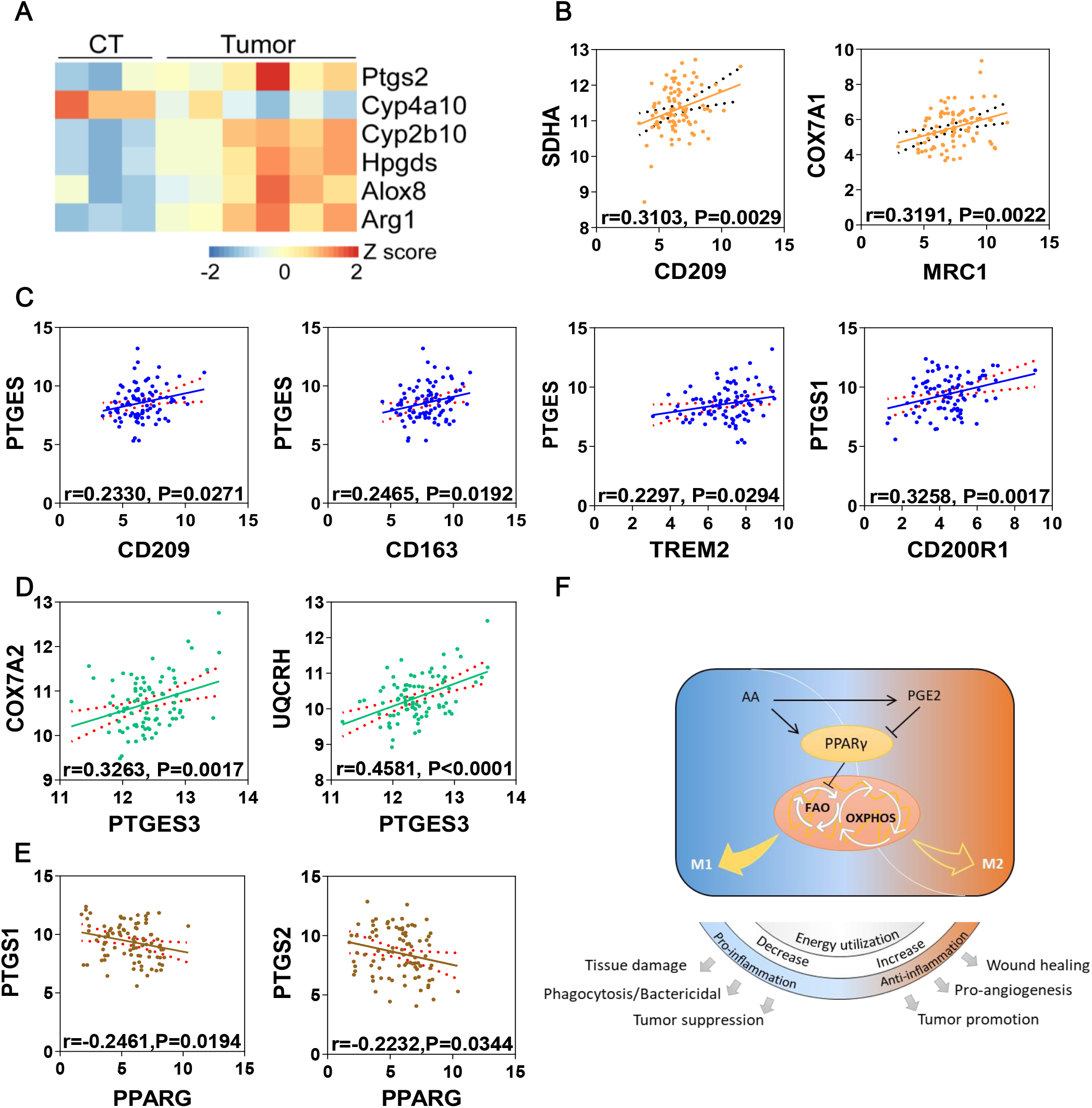
Arachidonic acid metabolism facilitates M2 TAM formation in ESCC. (A) Heatmap of differentially expressed genes matching “arachidonic acid metabolism” from mice ESCC or corresponding control tissues (CT). Arg1 was used as M2 macrophage marker. (B) Correlation analysis of oxidative phosphorylation (COX7A1, SDHA) and M2 macrophages (CD209, MRC1) in human ESCC from TCGA database. (C) Correlation analysis of PGE2 biosynthesis (PTGES, PTGS1) and M2 macrophages (CD209, CD163, CD200R1, TREM2) in human ESCC from TCGA database. (D) Correlation analysis of oxidative phosphorylation (COX7A2, UQCRH) and PGE2 biosynthesis (PTGES3) in human ESCC from TCGA database. (E) Correlation analysis of PPARG and PGE2 biosynthesis (PTGS1, PTGS2) in human ESCC from TCGA database. (F) Schematic diagram of arachidonic acid metabolism in controlling macrophage polarization and biological processes.

## Discussion

Distinct metabolic characteristic help macrophages with particular function during phenotype polarization (Saha et al., 2017). Lipid mediators are key fatty acid metabolites molecules involved in this process (Saha et al., 2017), serving as important signals. In the present study, we demonstrate that arachidonic acid metabolism is up-regulated upon M2 macrophage polarization. AA supplement inhibits IL-4/IL-13 stimulated M2 polarization of macrophages. PGE2, an essential metabolite generated from arachidonic acid metabolism, promotes macrophage polarization through inhibiting PPARG. Contrary to PGE2, inhibiting arachidonic acid metabolism to reduce PGE2 production suppresses M2 macrophage polarization. Our data elucidates a previously unappreciated mechanism of AA derived PGE2 to regulate macrophage alternative activation through directly de-activating PPARG, which de-activation indirectly facilitates OXPHOS by controlling FAO pathway. The newly uncovered connection between arachidonic acid metabolism and macrophage alternative activation is briefly outlined in Fig 6F. These metabolism controlled macrophages polarization will be finally reflected in many physiological and pathological processes.

Metabolism intricately links to immune homeostasis, which largely reflects by polarization of immune cells such as macrophages and T cells towards differential phenotypes. Like any physiological process, polarization of immune cells requires energy as well as the availability of nutrients, metabolites and oxygen (O’Neill et al., 2016). Macrophages in a nutrient deprivation or hypoxia microenvironment acquire distinct phenotype with those in perivascular areas (Laoui et al., 2014), suggesting an essential role of oxidative metabolism of nutrients. Actually, OXPHOS in mitochondria has been widely accepted as characteristic and necessity for M2 macrophages (Huang et al., 2014, Vats et al., 2006). This biological process is essential in M2 macrophages for ATP and biosynthetic output. Roisin *et al* has reviewed how oxidative metabolism control immune cell function (Loftus and Finlay, 2016). Inhibition of OXPHOS by reducing substrates or inhibiting mitochondrial complex had been found to suppress M2-related gene (Arg1, Mrc1) and surface marker (CD206) (Vats et al., 2006, Van den Bossche et al., 2016). In line with these observations, we found M2 surface marker (CD209) was suppressed by specific OXPHOS inhibitors in dose-dependent manner, suggesting that OXPHOS directly control macrophage alternative activation (M2 polarization).

Mitochondria utilize pyruvate from glucose metabolism, α-KG from glutamine metabolism as well as FAO derived acetyl-CoA to feed Krebs cycle and drive OXPHOS. It has been known that in M2 macrophages, FAO fuels OXPHOS thus provides a crucial energy source for M2 polarization (Vats et al., 2006, Huang et al., 2014). Blocking FAO pharmaceutically usually diminishes immune function of M2 macrophages or dampens M2 polarization (Huang et al., 2014) and favors M2-to-M1 repolarization (Hossain et al., 2015), suggesting an essential role of FAO in M2 macrophage polarization. We also verify its contribution on polarization by FAO inhibitor, which significantly decreased CD209 dose-dependently. Due to PPARs, especially PPARG, have been extensively investigated as essential nuclear receptors for M2 macrophage polarization and in lL-4 stimulated M2 polarization, PPARG and PGC1β are well known regulator of FAO (Odegaard et al., 2007, Vats et al., 2006), we speculated that PPARG regulated FAO might have a role in M2 polarization. Interestingly, our data revealed that PPARG negatively regulated CPT1A (an important enzyme for FAO) during M2 polarization, suggesting an inhibition of FAO by PPARG. Collectively, these data demonstrate that FAO regulated by PPARG are involved in M2 polarization.

In our study, PGE2 was found to dramatically promote M2 polarization and enhanced OXPHOS during this polarization process, revealing a possibility that PGE2 promotes M2 macrophage polarization through enhancing OXPHOS. On the basis of previous studies, PPARG regulated FAO may involve in this process (Namgaladze and Brune, 2016). Besides, our results show that PGE2 directly suppress PPARG expression and transcription activity (Fig S6). Therefore, it’s conceivable that inhibition of PPARG to favor FAO can enhance OXPHOS and lead to the observed promotion of PGE2 in M2 polarization. Consistently, activating PPARG might be a potential mechanism of AA inhibiting M2 polarization. In addition, PGE2-EP4 signaling has been reviewed as a possible mechanism to M2 polarization (Take et al., 2020), while in our polarization model, we found PGE2 increased M2 polarization even though EP4 was blocked (Fig S7). This suggests the existence of EP4-independent mechanisms under PGE2 mediated M2 polarization. Combining with the findings related to PPARG, we believe this PPARG dependent mechanism could explain large part of PGE2 mediated M2 polarization.

The identification of PGE2 as a key player for M2 macrophage polarization adds a metabolic explanation for how tumor polarize infiltrated macrophage towards an immunosuppressive M2 type. We found that several key AA metabolism associated genes were increased in mice ESCC, which was in line with increased M2 macrophage (indicated by Arg1), suggesting existence of a correlation between AA metabolism and M2 macrophage polarization in ESCC. Correlation analysis from human ESCC also demonstrates similar correlation between PGE2 biosynthesis and M2 macrophage polarization. These findings remind us that AA metabolizing towards PGE2 may be a potential prevention target for many M2 macrophages related diseases, such as cancers. Reducing PGE2 production by pharmacologically inhibiting COX-1/COX-2 might be a good strategy for cancer prevention. Actually, INDO, a non-specific COX-1/COX-2 inhibitor, has shown chemo-preventive and chemotherapeutic efficacy on colorectal cancer (Hull et al., 2003). In addition, it should be note that large amount of fatty acids and lipid mediators are pan-agonist for PPARs, thus PPARα and PPARδ may also involve in AA metabolism regulated M2 polarization. There is a need to clearly compare polarization effects between PPARs. Besides, many other metabolites could also be produced by AA metabolism, the polarization effect of this pathway may be more complex beyond our data suggest. Further investigations are needed to clearly clarify our findings.

In summary, we identified that arachidonic acid metabolism notably impact IL-4/IL-13 stimulated M2 macrophage polarization through regulating PPARG and OXPHOS. As one of the major metabolite of AA, PGE2 plays a crucial role in promoting macrophage polarization by the inhibition of PPARG and elevation of OXPHOS. Our finding renews the current understanding about the function roles of arachidonic acid metabolism as an immune regulator.

## Materials and Methods

### Reagents and antibodies

Arachidonic acid with purity > 98.5%, PGE2 and phorbol 12-myristate 13-acetate were obtained from Sigma-Aldrich (St. Louis, MO, USA). Cytokines (IL-4, IL-13, INFγ) were provided by Peprotech (Cranbury, NJ, USA). Fluorescence labeled antibodies (CD209,CD206 *et al*) and multi cytokines kit were from Biolegend (San Diego, CA, USA). Specific inhibitors were acquired from MedChemExpress (Monmouth Junction, NJ, USA) and Selleckchem (Houston, TX, USA). More information were offered in Supplementary table S1.

### Humanized THP-1 derived macrophage polarization model

Human monocytic THP-1 were obtained from American Type Culture Collection. THP-1 or their derived macrophages were maintained in RPMI-1640 medium supplemented with 10% fetal bovine serum, 1% penicillin-streptomycin and 0.05 Mm 2-mercaptoethanol at atmosphere of 37°C, 95% humidity, 5% CO_2_. To acquire macrophage, THP-1 were treated with phorbol 12-myristate 13-acetate (PMA, 25ng/ml) for 2 days and rest in PMA-free growth medium for one day to obtain undifferentiated macrophages (M0). Interferon-gamma (INF-γ, 25ng/ml) and Lipopolysaccharide (LPS, 100ng/ml) were added into M0 for another one day to obtain classically activated macrophage (M1); Interleukin-4 (IL-4, 20ng/ml) and Interleukin-13 (IL-13, 20ng/ml) treated M0 for 24 hours lead to M2 polarized macrophages. During macrophage polarization, specific agonists, inhibitors or arachidonic acid, PGE2 were co treated with cytokines. Anti-CCR7 and anti-CD209 fluorescence labeled antibodies were introduced to validate M1/M2 macrophages with high content screening system, respectively. Cell supernatants were collected for further analysis.

### Mice bone marrow derived monocytes based macrophage polarization model

Wild type C57BL/6 mice were obtained from Beijing Vital River Laboratory and C57BL/6 PPARγ^loxP^ mice (Pparg^tm2Rev/J^) were obtained from the Jackson laboratory. Monocyte-specific PPARG deletion mice (Pparg^-/-ΔMono^) were generated by breeding Pparg^tm2Rev/J^ with Lyz2^cre^ mice. Tibiae and femurs were isolated from 12 weeks old mice. Bone marrow medium (BMM) were prepared with DMEM containing 10% fetal bovine serum, 1% penicillin/streptomycin and 10 ng/ml M-CSF. Using syringe containing BMM with suitable needle to flushing out bone marrow. Next, removing debris or any remnants with strainers, centrifuged to obtain bone marrow derived cells. Re-suspended these cells in BMM and cultured for 7 days. BMM was refreshed on day 3, day 5. On day 7, collected macrophages were validated through flow cytometry and re-plated into multi-well plates for further test. For polarization, added INF-γ (25 ng/ml) plus LPS (100ng/ml) with/without treatments for M1 polarization, IL-4 (10 ng/ml) plus IL-13 (10 ng/ml) with/without treatments for M2 polarization. After 48h, macrophage markers were detected by high content screening system.

### Mice esophageal squamous cancer carcinoma (ESCC) model

This model was established as previously described (Yang et al., 2018). Briefly, N-Nitrosodimethylamine (NMBA) were administered to C57BL/6 mice by gavage at the dose of 0.25 mg/kg BW, twice weekly for 5 weeks. The control group (CT) was given the solvent Carboxyl Methyl Cellulose (CMC, 1%) with equivalent volume. All mice were housed in controlled atmosphere with 12h/12h light/dark cycle and fed with standard chow diet. After gavage, mice were observed for 20 weeks. At the endpoint of experiment, mice were sacrificed and forestomach of mice were collected and flash frozen in liquid nitrogen for RNA-seq. Transcriptomics data are available in GEO database (http://www.ncbi.nlm.nih.gov/geo/) under the accession number GSE134067.

### High content screening system (HCSS) for protein expression

For surface marker staining, cells in black wall 96-well-plate were washed with PBS for once and blocked with FcR blocking buffer (FcX block, Biolegend, San Diego, CA, USA) at room temperature for 10 minutes. Afterwards, replaced blocking buffer with cell staining buffer containing antibodies and Hoechst 33342 for another 30 minutes at room temperature. Next, washed with PBS and replaced with FluoroBrite ™ DMEM (Gibco, Grand Island, NY, USA). For intracellular protein staining, cells were fixed with fix/perm buffer (BD Bioscience) for half an hour and wash with Perm/Wash buffer (BD Bioscience) for two times prior to primary antibodies incubation. Then primary antibodies conjugated at room temperature for 30 minutes. After washing with Perm/Wash buffer, fluorescence conjugated secondary antibodies were incubated for another 30 minutes. Finally, stained for nucleic with Hoechst 33342 and acquired imagine and analysis with ImageXpress software (Molecular Device).

### RNA-seq and GSEA analysis

Total RNA was extracted from cells or tissues with RNeasy kit (QIAGEN) or TRIzol. RNA quality and quantity was detected with NanoDrop and Agilent 2100 Bioanalyzer. After passing the sample test, mRNA was enriched with magnetic beads with Oligo (dT) and broke into short fragments for cDNA synthesis. The cleaved RNA fragments were reversely transcribed into first strand cDNA using random hexamers, following by second strand cDNA synthesis using DNA Polymerase I and RNase H. The double-stranded cDNA was purified and added A tail and connected with a sequencing adapter. Then, perform PCR amplification on ABI StepOnePlus Real-Time PCR System and use the constructed sequencing library to sequence at Illumina HiSeq. Raw RNA sequencing data is available through the National Center for Biotechnology Information Gene Expression Omnibus (NCBI-GEO) database (http://www.ncbi.nlm.nih.gov/geo/) under the accession number GSE159112, GSE159120.

Raw data was filtered and clean reads were aligned with reference genome (hg19) using HISAT. Total mapped reads of all samples are higher than 95%. Reads were reconstructed into transcripts and their abundance was estimated and expressed as FPKM. DEseq2 was used to determine DEGs and |log2 (fold change)| ≥ 1 and adjust P value ≤ 0.05 were selected as significant difference. KEGG pathway analysis was performed with R phyper. Heatmap were generated with online tools (http://www.ehbio.com/ImageGP/index.php/Home/Index).

All FPKM value of identified genes from RNA-seq were input into GSEA software 4.0.3 for enrichment analysis (Subramanian et al., 2005). Data were normalized first and then a ranked gene list were generated. Database were downloaded from Molecular Signatures Database (MSigDB) gene sets (http://software.broadinstitute.org/gsea/index.jsp).

### Lipid metabolomics sample preparation and analysis

Differentiated and polarized M1/M2 were collected and immediately stored at liquid nitrogen until analysis. Thawed sample on ice and added 800 μl pre-chilled dichloromethane / methanol (3: 1) buffer, then precipitated in refrigerator at −20 *°C* for 2 hours. Then centrifuged at 25,000g, 4 °C for 15 minutes and transferred the supernatant 650μL to a new EP tube and centrifuged again. Then took 600 μL of supernatant to freeze-dry and added 600 μL lipid reconstituted solution (isopropanol: acetonitrile: water = 2: 1: 1) for reconstitution. After centrifuging, took 60 μL supernatant and 20 μL mixed QC for each sample to test on the LC-MS system.

Raw data from mass spectrometer were firstly preprocessed (noise filtering, peak matching and extraction), and corrected based on the Quality control-based robust LOESS signal correction (QC-RSC). Human Metabolome Database (HMDB) and LipidMaps database were used for peak alignment. Secondly, multivariate analysis PCA and PLS-DA were introduced to find out the differentially expressed metabolites. Metabolites with fold change ≥ 1.2 or ≤ 0.8333 and q-value < 0.05 were marked as significant difference. Finally, the differential metabolites identification and pathway analysis was performed. Data process and ion identification were performed with Progenesis QI (version 2.2) software. Pathway analysis was based on KEGG database.

### MSEA and Joint pathway analysis

Differential lipid metabolites from positive of negative ion mode were input for Metabolites Set Enrichment Analysis with online tools (MetaboAnalyst, https://www.metaboanalyst.ca) as previous study introduced (Chong et al., 2018). Differential lipid metabolites and differentially expressed genes from RNA-seq were input simultaneously to conduct joint pathway analysis on MetaboAnalyst. Integrated metabolic pathway database from current KEGG version were chose for enrichment. Parameter listed as follow: hyper geometric test for enrichment analysis, closeness centrality for topology measure and overall combine p value for integration method.

### Macrophage cytokines determination

After polarization with or without compounds of interest, medium containing treatments were replaced with fresh growth medium. 24 hours later, cell supernatants were collected for cytokines determination. The alternation of released cytokine from M1/M2 were detected with LEGENDplex™ Human macrophage panel using flow cytometry following manufacturer’s instruction. Data were analyzed with Legendplex software.

### Correlation analysis within TCGA data

Correlation analysis data used in this study is available in the Genomic Data Commons (https://portal.gdc.cancer.gov/). Briefly, a total of 90 cases of ESCC were included, and clinical characteristics had been supplied in previous study (Yang et al., 2020). Gene expression FPKM value were transformed with log2(X+1) (X= raw FPKM) for analysis. Pearson correlation coefficients and liner regression were computed within Graphpad Prism 6.

### Statistical analysis

All quantitative experiment values were expressed as mean ± SEM. Data were processed and visualized within Graphpad Prism 6. Unpaired t test or ANOVA analysis were applied to determine statistical significance within different treatments. P < 0.05 was set for significance.

## Supplementary Information

Supplementary figures and tables

## Acknowledgement

This work was supported by National Natural Science Foundation of China (No.81773437).

