## Supplementary figures and tables for "Arachidonic acid metabolism controls macrophage alternative activation through regulating oxidative phosphorylation in PPARG dependent manner"

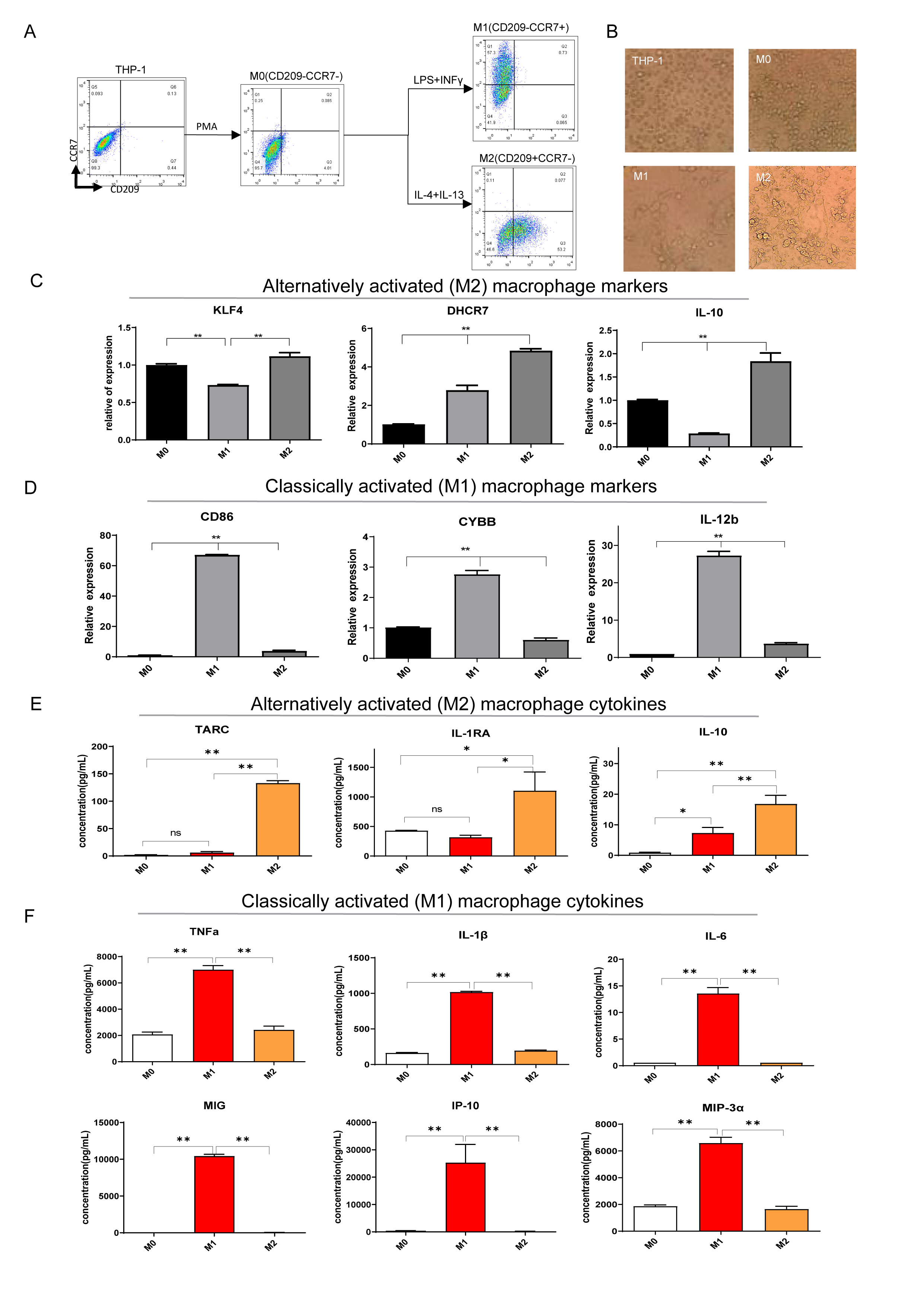

**Figure S1	Multiple parameters validated characteristics of THP-1 derived M1 M2.** (A) Surface marker (CCR7+CD209) detected by flow cytometry distinguished M1/M2. (B) Morphology alternation during differentiation and polarization. (C and D) qRT-PCR detected M2 or M1 macrophage markers. (E and F) Functional cytokines of M2 or M1 detected by legendplex panel. Error bars represent the mean ± SD from 3 biological replicates. *P< 0.05; ** P < 0.01.

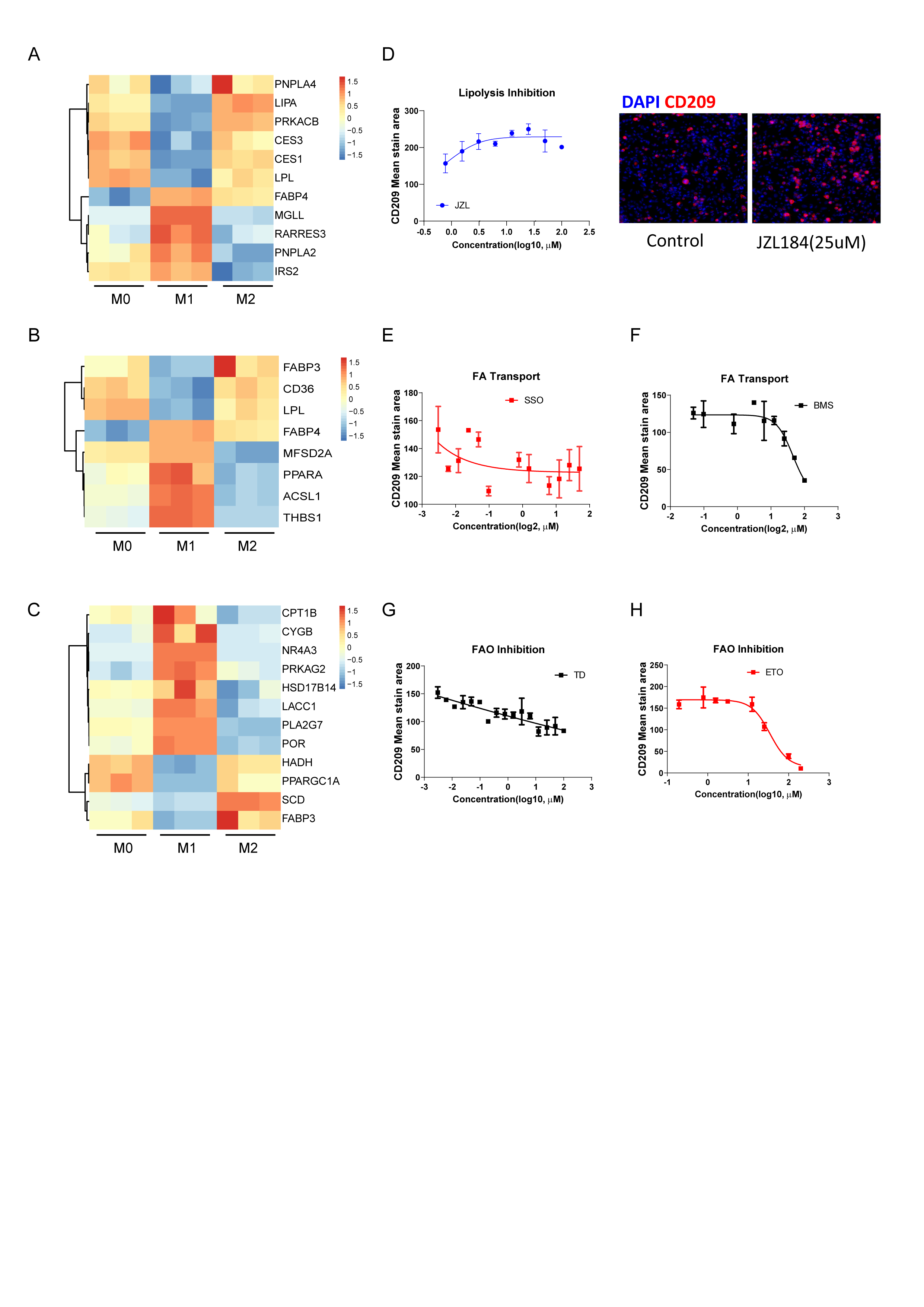

**Figure S2** **Lipid metabolism closely correlate to macrophage polarization.** (A-C) Gene expression in THP-1 derived differentially activated macrophage related to lipolysis, fatty acid transport or fatty acid oxidation, respectively. (D-H) Analysis of CD209 expression of THP-1 derived M2 macrophages via HCSS with inhibitors treated as indicated. Error bars represent the mean ± SD from 3 biological replicates. Right panel of (D): representative images.

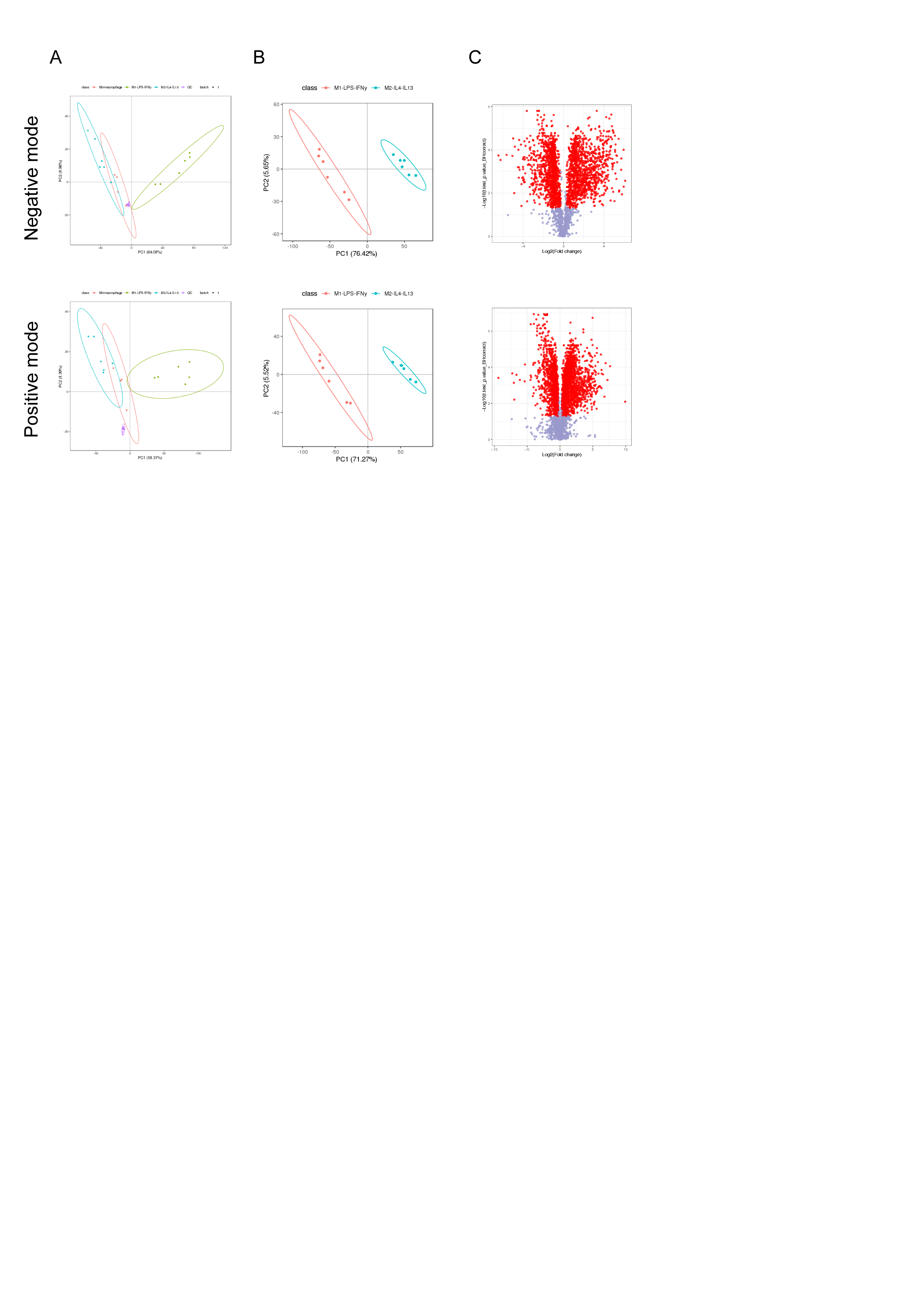

**Figure S3	Distinct lipidomics characteristics of M1/M2 macrophages**. (A) PCA including quality control samples QC indicated a good quality of the collected data. (B) PLS-DA revealed distinct metabolites profile of M1/M2 macrophages. (C) Overall differential metabolites of M1/M2 macrophages. Fold change ≥ 1.2 or ≤ 0.83 and q-value < 0.05 were set as screening condition. Red: differential metabolites; grey: no difference metabolites.

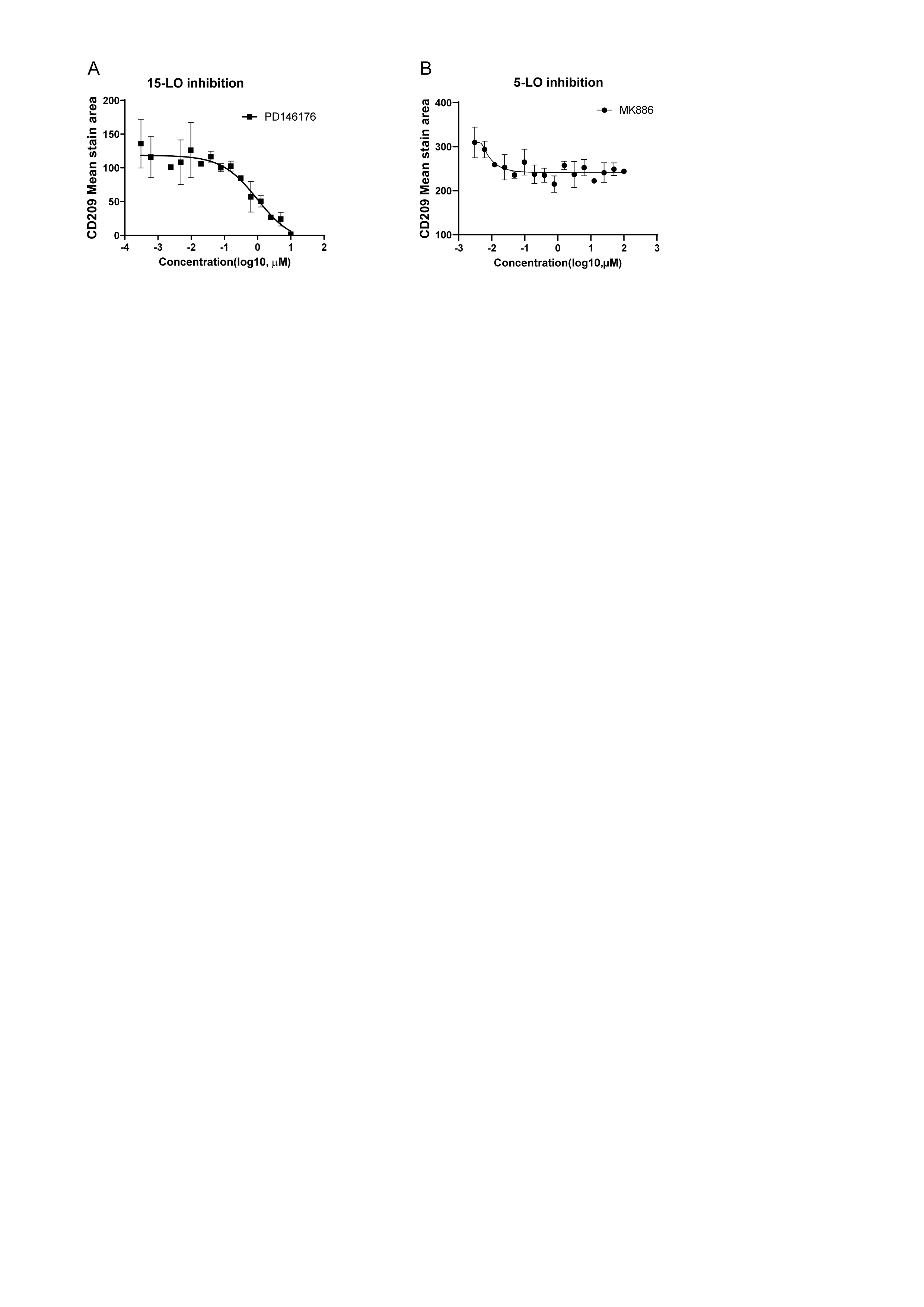

**Figure S4	Lipoxygenases inhibition in arachidonic acid metabolism decreased M2 macrophages polarization dose dependently.** (A and B) Dose response curve for 15-lipoxygenase (15-LO) or 5-lipoxygenase (5-LO) in M2 polarization. Error bars represent the mean ± SD from 3 biological replicates.

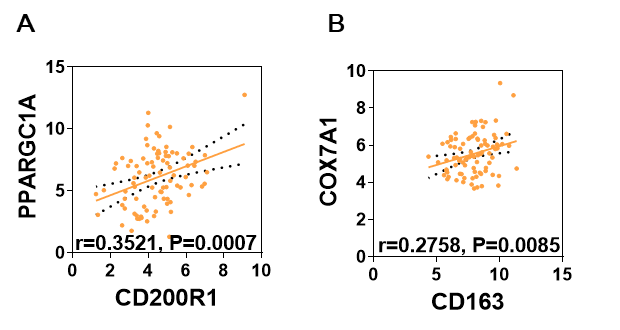

**Figure S5	Oxidative phosphorylation correlates to M2 TAM in ESCC.**

(A and B) Correlation analysis of oxidative phosphorylation (PPARGC1A, COX7A1) and M2 macrophages (CD200R1, CD163) in human ESCC from TCGA database.

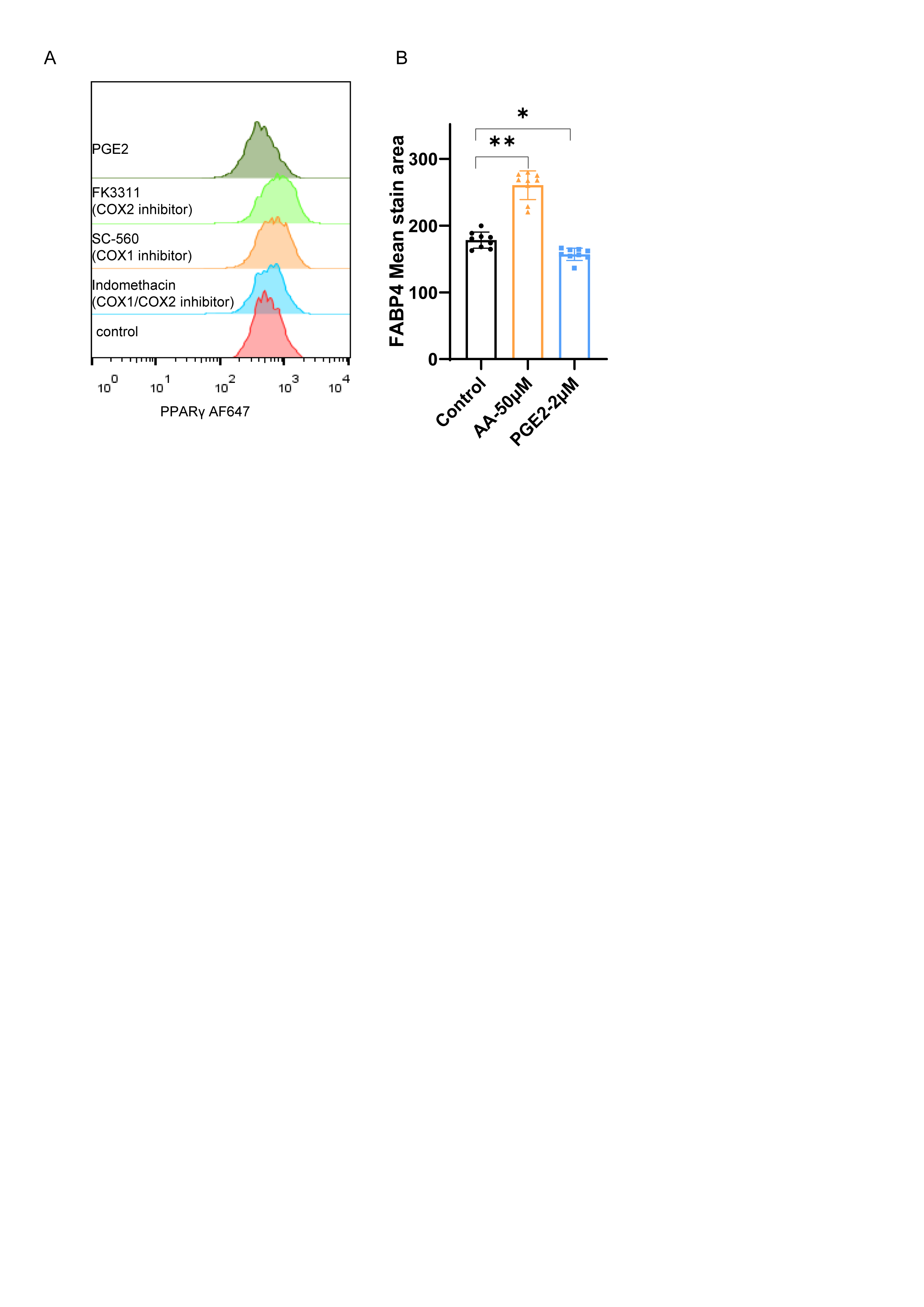

**Figure S6	Expression and transcription activity of PPARG were inhibited by PGE2 treatment.** (A) Representative fluorescence histogram of macrophages PPARG from flow cytometry with treatment as indicated. (B) Expression of macrophages FABP4 (indicator for PPARG transcription activity) from HCSS with treatment as indicated. Error bars represent the mean ± SD from 3 biological replicates. *P< 0.05; ** P < 0.01.

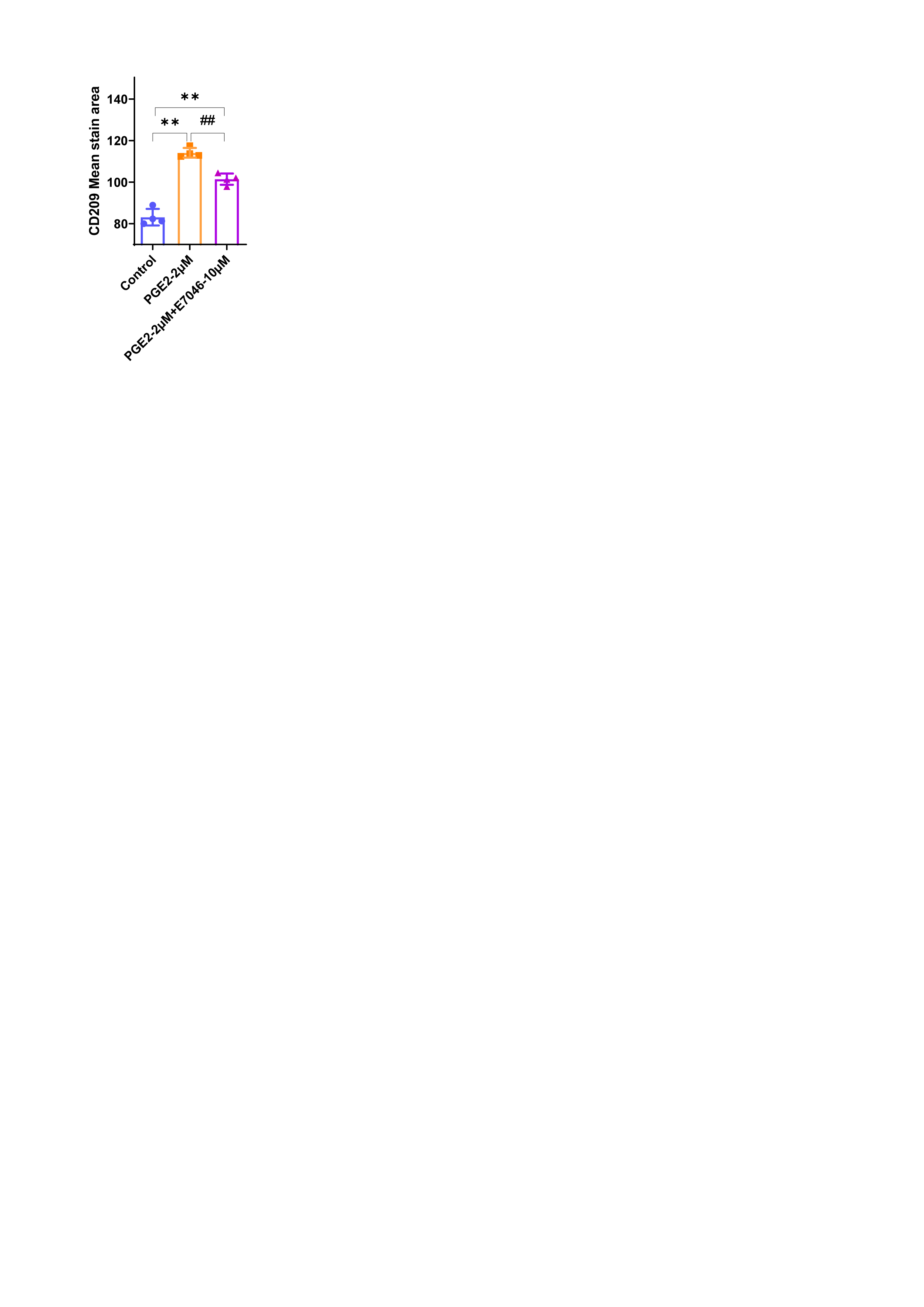

**Figure S7	Blockade of EP4 did not abolish PGE2 promoted M2 polarization.** Error bars represent the mean ± SD from 3 biological replicates. Compared with control, ** P < 0.01. Compared with first treatment, ^##^ P < 0.01.

**Table S1 Key resources**

| **REAGENTS OR RESOURCES** | **SOURCE** | **IDENTIFIER** |
| --- | --- | --- |
| **Biological material** |  |  |
| THP-1 | American Type Culture Collection | TIB-202 |
| Bone marrow derived monocytes (wild type) | C57BL/6 mice | / |
| Bone marrow derived monocytes (PPARG KO) | Jackson Labs | / |
| **Chemicals, Antibodies, cytokines** |  |  |
| Anti-CCR7 | Biolegend | Cat#353204 |
| Anti-CD209 | Biolegend | 330106;330104 |
| Anti-CD206 | Biolegend | 141706 |
| Anti-CD69 | Biolegend | 104518 |
| Arachidonic acid | Sigma Aldrich | A3611 |
| ATP5A | Abcam | ab14748 |
| BMS309403 | MCE | HY-101903 |
| CPT1A | Proteintech | 15184-1-AP |
| Etomoxir sodium salt | MedChemExpress | HY-50202A |
| FABP4 PE | LSBio | LS-C650165 |
| Human Trustain FcX | Biolegend | 422302 |
| Fatostatin | MedChemExpress | HY-14452 |
| FK 3311 | MedChemExpress | HY-14445 |
| FT113 | MedChemExpress | HY-111551 |
| Human IL-4 | Peprotech | 200-04 |
| Human IL-13 | Peprotech | AF-200-13 |
| Human IFNγ | Peprotech | 300-02 |
| IACS-10759 | Selleck | S8731 |
| Indomethacin | MedChemExpress | HY-14397 |
| JZL184 | MedChemExpress | HY-15249 |
| Lipopolysaccharides (LPS) | Sigma Aldrich | L6529 |
| Murine M-CSF | Peprotech | 315-02 |
| Murine IL-4 | Peprotech | 214-14 |
| Murine IL-13 | Peprotech | 210-13 |
| Murine IFNγ | Peprotech | 315-05 |
| 3-Nitropropionic acid | Selleck | S3652 |
| PPARgamma | Cell signaling technology | 2435S |
| Prostaglandin E2 | Sigma Aldrich | P0409 |
| PMA | Sigma Aldrich | P8139 |
| Sulfosuccinimidyl oleate(SSO) | Sigma Aldrich | SML2148 |
| VLX600 | MedChemExpress | HY-12406 |
| **Critical Commercial assays** |  |  |
| Foxp3 / Transcription Factor Staining Buffer Set | Invitrogen | 00-5523-00 |
| LEGENDplex Human Macrophage/Microglia Panel | biolegend | 740502 |
| RNeasy Mini Kit | QIAGEN | 74104 |
| **Deposited Data** |  |  |
| RNA-sequencing data | This paper; previous study ^[1]^ | GSE159112, GSE159120; GSE134067 |
| **Software and Algorithms** |  |  |
| Gene Set Enrichment Analysis (GSEA) (v4.0.3) | ^[2]^ | http://software.broadinstitute.org/gsea/index.jsp |
| Metabolite Set Enrichment Analysis (MSEA)(v4.0) | ^[3]^ | https://www.metaboanalyst.ca/ |
| Joint Pathway analysis | ^[3]^ | https://www.metaboanalyst.ca/ |
| GraphPad prism (v 6.01) | GraphPad Software | N/A |
| ImageXpress | Molecular Device |  |
| **Other** |  |  |
| DMEM | Gibco | 11995065 |
| Fetal Bovine Serum (FBS) | Gibco | 10099141C |
| Fixation and Permeabilization Solution | BD Bioscience | 554722 |
| FluoroBrite ^TM^ DMEM | Gibco | A1896702 |
| Hoechst 33342 | Solarbio | C0031 |
| penicillin-streptomycin | Gibco | 15140122 |
| Perm/Wash | BD Bioscience | 554723 |
| RPMI 1640 (ATCC modification) | Gibco | A1049101 |
| 2-mercaptoethanol | Sigma Aldrich | M3148 |

Table S2 Arachidonic acid metabolism associated metabolites from lipidomics analysis

| **mode** | **ratio** | **Lable** | **description** |
| --- | --- | --- | --- |
| negative | 4.06 | 6.58_397.2261m/z | Prostaglandins(PGH2;PGE2;PGD2;PGI2);TXA2;Lipoxin A4;Lipoxin B4;20-hydroxy LTB4;15-keto-PGF2alpha |
| positive | 2.82 | 0.96_372.2761m/z | 11beta-PGF2;15-F2t-IsoP;PGF2alpha;11,12,15-THETA;Trioxilin A3;11,14,15-THETA |
| positive | 2 | 1.43_586.3112m/z | LTF4 |
| negative | 1.76 | 3.04_411.2041m/z | 20-carboxy-LTB4 |
| positive | 1.29 | 3.05_327.2306m/z | Arachidonic acid |
| positive | 0.78 | 1.12_338.2474n | 5,6-DHET;8,9-DiHETrE;11,12-DiHETrE;14,15-DiHETrE |
| positive | 0.75 | 0.76_334.2349m/z | 15-Deoxy-d-12,14-PGJ2 |
| positive | 0.72 | 1.13_317.2101m/z | 12-oxo-LTB4;PGA2;PGB2;PGC2;PGJ2;delta-12-PGJ2;5,6-Ep-15S-HETE |
| positive | 0.65 | 2.87_336.2517m/z | LTA4;15-KETE;5-KETE;12-KETE |
| negative | 0.41 | 0.68_355.2036m/z | 15S-HETE;5S-HETE;20-HETE;19S-HETE;5,6-EET;(+/-)8,9-EpETrE;(+/-)11,12-EpETrE;(+/-)14,15-EpETrE;8S-HETE;12S-HETE;16R-HETE;9S-HETE;11R-HETE;12R-HETE;8R-HETE |

Table S3 Detail information for top 20 GSEA enrichment pathways in PGE2 treated M2 polarization.

| **NAME** | **SIZE** | **ES** | **NES** | **NOM p-val** | **FDR q-val** |
| --- | --- | --- | --- | --- | --- |
| STAPHYLOCOCCUS_AUREUS_INFECTION_-_HOMO_SAPIENS_(HUMAN)(HSA05150) | 301 | 0.66548 | 1.40566 | 0 | 0.738042 |
| ANTIGEN_PROCESSING_AND_PRESENTATION_-_HOMO_SAPIENS_(HUMAN)(HSA04612) | 211 | 0.722176 | 1.392314 | 0 | 0.570246 |
| IL-17_SIGNALING_PATHWAY_-_HOMO_SAPIENS_(HUMAN)(HSA04657) | 140 | 0.510845 | 1.376146 | 0 | 0.489651 |
| PHAGOSOME_-_HOMO_SAPIENS_(HUMAN)(HSA04145) | 453 | 0.509968 | 1.357369 | 0 | 0.465207 |
| OSTEOCLAST_DIFFERENTIATION_-_HOMO_SAPIENS_(HUMAN)(HSA04380) | 191 | 0.461932 | 1.343462 | 0 | 0.443978 |
| HEMATOPOIETIC_CELL_LINEAGE_-_HOMO_SAPIENS_(HUMAN)(HSA04640) | 199 | 0.655428 | 1.341753 | 0 | 0.386636 |
| TH17_CELL_DIFFERENTIATION_-_HOMO_SAPIENS_(HUMAN)(HSA04659) | 166 | 0.533104 | 1.338639 | 0 | 0.338688 |
| GLYCOSAMINOGLYCAN_BIOSYNTHESIS_-_CHONDROITIN_SULFATE_/_DERMATAN_SULFATE_-_HOMO_SAPIENS_(HUMAN)(HSA00532) | 28 | 0.601301 | 1.332624 | 0.181818 | 0.317932 |
| TUBERCULOSIS_-_HOMO_SAPIENS_(HUMAN)(HSA05152) | 398 | 0.540087 | 1.332295 | 0 | 0.288273 |
| LEISHMANIASIS_-_HOMO_SAPIENS_(HUMAN)(HSA05140) | 242 | 0.591083 | 1.328794 | 0 | 0.267774 |
| OXIDATIVE_PHOSPHORYLATION_-_HOMO_SAPIENS_(HUMAN)(HSA00190) | 861 | 0.543396 | 1.32836 | 0 | 0.248068 |
| INFLAMMATORY_BOWEL_DISEASE_-_HOMO_SAPIENS_(HUMAN)(HSA05321) | 100 | 0.584769 | 1.326967 | 0 | 0.231645 |
| VARIOUS_TYPES_OF_N-GLYCAN_BIOSYNTHESIS_-_HOMO_SAPIENS_(HUMAN)(HSA00513) | 66 | 0.573851 | 1.310889 | 0 | 0.240281 |
| GRAFT-VERSUS-HOST_DISEASE_-_HOMO_SAPIENS_(HUMAN)(HSA05332) | 99 | 0.752296 | 1.30874 | 0 | 0.226761 |
| TYPE_I_DIABETES_MELLITUS_-_HOMO_SAPIENS_(HUMAN)(HSA04940) | 123 | 0.665351 | 1.305426 | 0 | 0.230762 |
| NON-ALCOHOLIC_FATTY_LIVER_DISEASE_-_HOMO_SAPIENS_(HUMAN)(HSA04932) | 626 | 0.494478 | 1.305105 | 0 | 0.219527 |
| TYROSINE_METABOLISM_-_HOMO_SAPIENS_(HUMAN)(HSA00350) | 67 | 0.48297 | 1.299975 | 0 | 0.227472 |
| VIRAL_MYOCARDITIS_-_HOMO_SAPIENS_(HUMAN)(HSA05416) | 276 | 0.578541 | 1.298735 | 0 | 0.217668 |
| ESTROGEN_SIGNALING_PATHWAY_-_HOMO_SAPIENS_(HUMAN)(HSA04915) | 322 | 0.459586 | 1.290178 | 0 | 0.222081 |
| RHEUMATOID_ARTHRITIS_-_HOMO_SAPIENS_(HUMAN)(HSA05323) | 209 | 0.583726 | 1.287354 | 0 | 0.217427 |
